# A dual-function variant on chromosome 17 regulates circRNA expression and splicing in multiple sclerosis

**DOI:** 10.64898/2026.03.18.712599

**Authors:** Saioa Gs Iñiguez, Leire Iparraguirre, Eduardo Andrés-León, Hirune Crespillo, Leire Romarate, Tamara Castillo Triviño, Elena Urcelay, Manuel Comabella, Sunny Malhotra, Xavier Montalban, Lluís Ramió-Torrentà, Anna Quiroga-Varela, Koen Vandenbroeck, Ane Aldekoa, Antonio Alcina, David Otaegui, Fuencisla Matesanz, Maider Muñoz-Culla

## Abstract

Multiple sclerosis (MS) is a chronic autoimmune demyelinating disease of the central nervous system with a complex etiology. Recent genomic studies highlight the contribution of expression quantitative trait loci (eQTLs) in modulating gene expression and disease susceptibility. Given the emerging role of circular RNAs (circRNAs) in MS, we hypothesized that genetic variants may regulate circRNA expression through circRNA-specific eQTLs (circ-eQTLs). We performed a *cis*-circ-eQTL analysis integrating circRNA expression and whole-genome genotyping data from 30 MS patients and 18 healthy controls using a linear regression model adjusted for disease status and sex. Candidate circ-eQTLs were prioritized based on MS-associated regions and known splicing QTLs (sQTLs) from GTEx and validated in an independent cohort (67 MS, 64 controls). Association analysis in a larger cohort (2831 MS, 3191 controls) evaluated two candidate variants for MS risk. We identified 42,077 significant *cis*-circ-eQTLs and validated three. Two SNPs, rs7214410 and rs11079784, modulated hsa_circ_0106983 expression, and rs7214410 also acted as an sQTL affecting *EFCAB13* splicing. rs7214410 showed stronger association with MS than rs11079784. Our findings reveal extensive genetic regulation of circRNA expression and highlight rs7214410 as a dual-function variant refining the MS susceptibility locus on chromosome 17.

## Introduction

Multiple sclerosis (MS) is a chronic autoimmune disorder that primarily affects the central nervous system (CNS) and is typically manifested in young adulthood, with an average age of diagnosis of 32 years old. The latest data estimate that a total of 2.8 million people suffer from MS worldwide and that the prevalence has increased in the last decade. Moreover, women are twice as likely to have MS as males, although this ratio is higher in some countries[1].

The etiology of MS is complex and multifactorial, where several factors such as environmental factors, genetics or microbiota take part as risk factors for developing MS[2]. The combined influence of these factors underlies the susceptibility of the disease and the understanding of their relationship, will help to unveil the causes behind the heterogeneity of people with MS (pwMS).

The pathophysiology of the disease is also complex and heterogeneous but many research articles in the last decades proved the involvement of the immune system in the pathology of MS. Evidence suggested that the disease involves an immune-mediated process where immune cells infiltrate the CNS and initiate an inflammatory response against myelin sheath that covers nerve fibers, leading to inflammation and demyelination[3]. Bidirectional interactions have also been shown among several immune cell subsets in the periphery (T cell, B cells, and myeloid cells) and resident CNS cells (microglia and astrocytes). All these cells secrete inflammatory molecules that recruit immune cells from the periphery to the CNS, generate an inflammatory response and lead to neuronal demyelination[4].

In this scenario, the understanding of molecular mechanisms that trigger and regulate immune response turns up crucial, and genetic and transcriptomic studies have been really enlightening to uncover molecular mechanisms involved in the immune function. In this line, transcriptomic studies have changed from a classical view of studying only the protein-coding fraction of the genome to analyzing the non-coding fraction as well. These studies have revealed that non-coding RNAs are also altered in the disease and contribute to the regulation of different cellular processes in the pathophysiology. Different non-coding RNA families have been identified to be differentially expressed in leukocytes from pwMS, such as miRNA, circular RNAs (circRNAs) or long non-coding RNAs, among others[5]–[9].

Besides, genome-wide association studies (GWAS) performed in the last two decades in big collaborative efforts have been successful in identifying genetic variants that confer a greater risk for developing MS, with over 200 autosomal susceptibility variants identified outside the major histocompatibility complex [10]–[13].

Functional annotation analyses have revealed that MS-associated variants are often located in or near genes related to immune function, suggesting a strong immunogenetic component[13]. In addition, it has been proposed that MS association variants may exert their effect by modulating the expression level of genes, acting as expression quantitative trait loci (eQTL). In fact, in the last genome-wide association study performed in MS, some eQTL have been identified in peripheral blood mononuclear cells, as well as in different immune cell subtypes and the prefrontal cortex[13]. Nonetheless, these analyses are usually focused on protein-coding genes, and they set aside non-protein coding genes. Other authors have demonstrated that genomic variants can act as eQTL for miRNA in humans[14]. In the context of disease, there are some works reporting that MS-associated variants can act as eQTL in blood cells for pri-miRNA and circRNA[15],[16], suggesting that the role of MS-associated variants in conferring a higher risk for developing MS can be due to differences in non-coding RNA expression.

Bearing in mind these concepts, the aim of this work is to study the effect of genetic variants on the expression of circRNA in leukocytes. To do so we have analyzed genome-wide variant associations with circRNA expression in peripheral leukocytes of pwMS and healthy controls (HC) and compare them with MS-associated regions. In this work, we demonstrate that the causal gene of the association with MS of a region in chromosome 17 can be a circRNA. Moreover, the variant causing the circ-eQTL is also responsible for a splicing QTL on the *EFCAB13* gene.

## METHODOLOGY

### Participants and blood Sampling

Whole blood was collected from a total of 97 people with MS (pwMS) and 84 healthy controls (HC) in the Department of Neurology at Donostia University Hospital. Participants in the MS group had received multiple sclerosis diagnosis according to the McDonald criteria and HC were individuals without any known neurological disease. The main clinical and demographic characteristics of both patients and healthy donors of these two cohorts are summarized in Table 1. They were under different treatments. Apart from that, they were not diagnosed with other autoimmune or infectious diseases.

**Table 1.** Main clinical and demographic characteristics of the individuals enrolled in the study.

|  |  | Type of MS | Sex | Age | EDSS | AOO |
| --- | --- | --- | --- | --- | --- | --- |
| Discovery Group | MS (n=30) | RR-MS (n=20) | Female (n=16) | 41.0 (± 10.6) | 3.0 (0.0-6.5) | 29.0 (± 6.8) |
| | | | Male (n=4) | 48.8 (± 7.4) | 3.4 (2.0-5.5) | 35.1 (± 8.4) |
| | | SP-MS (n=10) | Female (n=7) | 50.7 (± 3.9) | 5.3 (3.5-7.0) | 31.9 (± 8.6) |
| | | | Male (n=3) | 39.2 (± 7.4) | 6.2 (4.0-8.0) | 20.9 (± 5.4) |
| | HC (n=18) | - | Female (n=13) | 49.6 (± 7.9) | - | - |
| | | | Male (n=5) | 40.0 (± 10.9) | - | - |
| Validation group | MS (n=67) | PP-MS (n=3) | Female (n=3) | 65.0 (± 5.0) | 5.8 (4.0-7.5) | 46.7 (± 11.2) |
|  |  |  | - | - | - | - |
| | | RR-MS (n=63) | Female (n=47) | 43.1 (± 13.9) | 1.8 (0.0-7.0) | 29.4 (± 10.0) |
| | | | Male (n=16) | 42.8 (± 13.7) | 1.6 (0.0-4.0) | 29.6 (± 9.9) |
|  |  | SP-MS (n=1) | - | - | - | - |
|  |  |  | Male (n=1) | 38.0 | 4.5 | 18.0 |
| | HC (n=64) | - | Female (n=42) | 40.0 (± 18.8) | - | - |
| | | | Male (n=22) | 44.4 (± 19.8) | - | - |

Abbreviations: MS, multiple sclerosis; HC, healthy control; RR-MS, relapsing-remitting MS; SP-MS, secondary progressive MS; PP-MS, primary progressive MS; EDSS, Expanded Disability Status Scale; AOO, Age of Onset. Age and AOO data are presented as “average (standard deviation)”, and EDSS data are shown as “average (range)”.

An additional cohort including 2831 pwMS and 3191 HC was studied in a case-control association study. These participants were recruited from 6 different Spanish Hospitals: 870 pwMS and 1143 HC from Parasitology and Biomedicine López Neyra Institute-CSIC in Granada; 762 pwMS and 936 HC from Health Research Institute from San Carlos Clinic Hospital in Madrid; 356 pwMS and 321 HC from Recerca Institute from Vall d‘Hebron Hospital in Barcelona; 185 pwMS from Dr. Josep Trueta Biomedicine Research Institute in Girona; 347 pwMS and 591 HC from Biocruces-Bizkaia Health Research in Bilbao and 311 pwMS and 200 HC from Donostia University Hospital.

### RNA and DNA isolation

Samples were processed in the <u>Basque Biobank</u> to isolate total RNA and DNA following their normal procedures. RNA from all the cohorts of validation was isolated with the miRNeasy mini kit (Qiagen), following the manufacturer’s instructions. RNA concentration was measured using NanoDrop ND-1000 spectrophotometer (Thermo Scientific).

DNA from all the cohorts of validation was isolated in the Basque Biobank, with the Flexigene DNA kit (Qiagen) helped by the automatic AGFlex Star robot, following the manufacturer instructions and eluted in elution buffer TrisCl 10mM with pH= 8.5. DNA concentration was measured using NanoDrop ND-1000 spectrophotometer (Thermo Scientific). DNA samples from the association study cohort were obtained and analyzed in each center.

### CircRNA expression

CircRNA expression profile in peripheral blood leukocytes of the discovery cohort was measured by RNAseq and analyzed as described in a previous work[7][17]. The detection and quantification workflow in this cohort resulted in a list of 22,835 *bona fide* circRNA, which were included in the present study for *cis*-circ-eQTL mapping.

Selected candidate circ-eQTLs were validated in a larger and independent cohort of 68 pwMS and 64 HC using a two-step RT-qPCR approach. For validation experiments, total RNA was reverse transcribed into cDNA with random primers using High-Capacity cDNA Reverse Transcription Kit (Applied Biosystems, Inc., USA) according to the manufacturer’s instructions. The RT-PCR was performed in a Verity Thermal cycler (Applied Biosystems, Inc., USA) with the following program: 25°C for 10 min, 37°C for 120 min and 85°C for 5 min. Q-PCR was performed using Power SYBRGreen Master Mix (Applied Biosystems) in a CFX384 Touch Real-Time PCR Detection System (Bio-Rad Laboratories, Inc.). Each sample was run in triplicate (starting from 10 ng of cDNA in 10 μL total reaction volume). Q-PCR cycling conditions were: 95°C for 10 min, 40 cycles of 95°C for 15 s and 60°C 1 min followed by a dissociation curve analysis. The raw Cq values and melting curves were analyzed in CFX Maestro 1.0 (BioRad) where a single-peak in the melting curve indicated the specificity of the amplification.

CircRNA sequences were obtained from UCSC (University of California, Santa Cruz) Genome Browser (GRCh37/hg19 assembly) and divergent primers were designed to amplify the circular transcripts so that the qPCR amplicon spans the backspliced junction[18]. Primer3Plus software[19] was used to assist the primer design and to ensure the target specificity of the primers respectively. Primer sequences for hsa_circ_0106983 selected were Forward primer: 5`-AAATGAGAATGGAATGGTGGAG-3‘ and Reverse primer: 5`-ATCCAAAACAGCAAACACGTC-3‘, and for hsa_circ_00002161 selected ones are Forward primer: 5`-GGAGGACAAATAATTAGAAAATGGA -3‘ and Reverse primer: 5`-TTCCATCTCCCAAGGAACC -3‘. PCR amplicons were firstly subjected to Sanger Sequencing (ABIprism 3130), and checked for the presence of the predicted backspliced junction to test their circularity. In the case of the reference gene, we have used an already designed Beta-2-Microglobulin primers (Hs_B2M_1_SG Quantitect primer assay from Qiagen). The expression of circRNA was normalized against the reference gene and converted to the linear scale (2^-ΔCT^×1,000)

### Genotyping

Genome-wide genotyping of the samples from the discovery cohort was performed in the National Center of Genotyping from Spain (CEGEN) with Illumina Infinum Global Screening array. Genotyping of SNPs selected for validation was performed with TaqMan SNP Genotyping assays (Thermofisher Scientific) and TaqPath™ ProAmp™ Master Mix (A30865, Applied Biosystems, Foster City, CA, USA), in a Real-Time quantitative PCR system (Bio-Rad CFX384) following the manufacturer’s instructions. 10 ng of genomic DNA was used in each reaction. Allelic discrimination was performed with CFX Maestro v2.3 software and genotype assignments were confirmed by reviewing the RT-PCR spectra of each sample.

### *Cis*_Circ-eQTL mapping

First, an imputation step was carried out using only autosomal chromosomes. Quality control assessment of genome-wide genotyping results and imputation analysis resulted in 4,824,059 imputed variants. Circ-eQTLs analysis was performed correlating genotypes of 4,824,059 imputed variants in autosomal chromosomes with expression levels of 22,835 circRNAs. Only *cis-*eQTLs were considered (a maximum distance of 1Mb). A linear regression model was used controlling for group (MS or HC) and sex. Significant interactions were considered those with corrected p-value (FDR) < 0.05 and due to our discovery cohort size, an additional filtering threshold of minor allele frequency (MAF) > 0.1 was established.

### Selection of candidate circ-eQTL

The resulting circ-eQTLs were intersected with the 200 regions reported to be associated with MS susceptibility[13] to obtain candidate circ-eQTL for validation. Additionally, they were also intersected with splicing QTLs (sQTL) described in GTex database (https://gtexportal.org) (Fig. 1).

**Figure 1.**
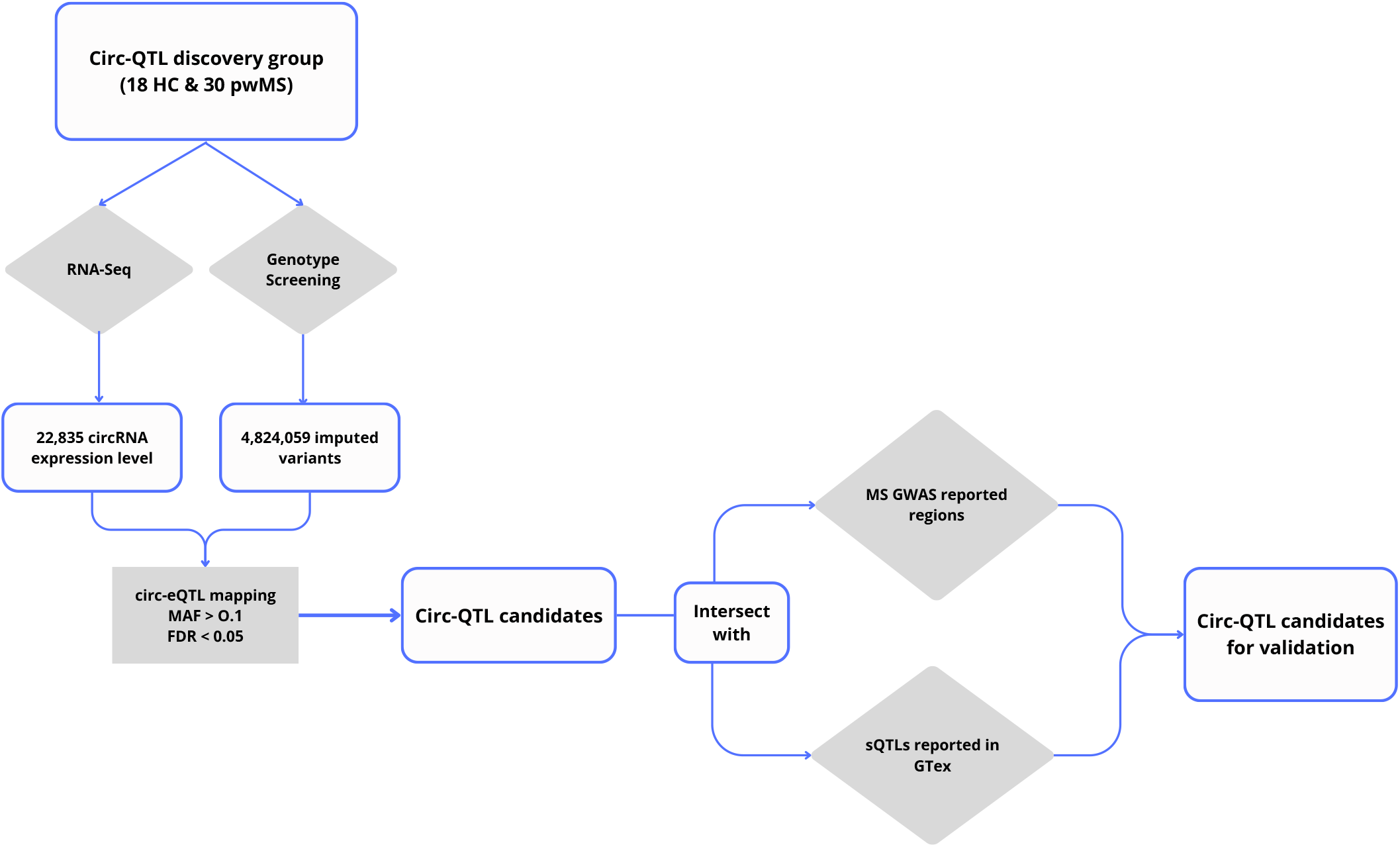
Circ-eQTL candidate selection workflow.

### Case-control association study

We performed an association study to see whether the SNP identified as a circ-eQTL was also associated with multiple sclerosis. The association study group is entirely comprised by 2831 pwMS and 3191 HC, with the collaboration of different research groups in Spain. First, linkage-disequilibrium (LD) was calculated between the SNP causing the circ-eQTL and the MS-associated SNP from the European Population using the LDmatrix tool, which is based on 1000 Genome data[20]. The aim of this analysis was to check whether we would be able to discriminate the effect of each SNP. Genotyping of these samples was performed in each lab using the same protocol as described in the previous section. Data were analysed using SNPassoc package version 2.1-0 in RStudio.

### Splicing validation study

We first used9 lymphoblastoid B-cell lines (LCL), selected 3 samples for each genotype and subsequently, 6 HC and 6 MS samples were selected for each genotype, from the validation group (18 HC and 18 MS samples). Both sample types followed a similar protocol. First, RNA was transcribed to cDNA using a reverse transcriptase enzyme that only amplifies linear transcripts to avoid the presence of circRNA fragments. We used the SuperScript IV Reverse Transcriptase enzyme (Thermo Fisher; Cat. No. 18090010), together with 10mM dNTPmix (Thermo Fisher, Cat. No. 18427-013), RNaseOUT Recombinant Ribonuclease Inhibitor (Thermo Fisher; Cat. No. 10777019) and the Oligo(dT)_20_ (Thermo Fisher; Cat. No. 18418-020) following the manufacturer’s instructions for the SuperScript IV First-Strand cDNA Synthesis Reaction kit, with a final volume of 20 μl and using 350ng of RNA sample as starting material.

The amplification of exons 8 - 11 of the *EFCAB13* gene was performed with previously designed forward primer (5`-GGAAAAGGAAATGCTGTCTAACC-3`), located in the start of exon 8, and reverse primer (5`-ATCCCCAATATCCACCATGTG-3`), located at the end of exon 11. The PCR was performed with Hot Start Taq 2x Master Mix kit (New England Biolabs, Massachusetts; Cat. No. M0496S) following the manufacturer’s instructions and adding 1 μl per reaction of 10 μM of each primer and 10 μl of 10ng/μl cDNA. The PCR cycling protocol was carried out in a Verity Thermal cycler (Applied Biosystems) as follows: 94°C for 30 sec and 30 cycles of 94°C for 15 sec, 59°C for 30 sec and 68°C for 35 sec, finally samples were kept at 68°C for 5 min.

Amplification product from LCL samples was loaded onto an acrylamide gel (12%) for the visualization of the sequences with pBR322 DNA/BsuRI (HaeIII) marker (Thermo Fisher Scientific). Amplification products of peripheral leukocyte samples were loaded on an agarose gel (3 %) with 1.5 μl of GelRed Nucleic Acid Gel Stain (Biotium) as a fluorescent nucleic acid dye, with ΦX174 DNA/HaeIII Marker. Results were analysed in the iBright Imaging System (Thermo Fisher Scientific) using ImageJ v1.54g software.

Finally, different size PCR amplicons were extracted from the agarose gel and purified by NucleoSpin gel and PCR Clean-up mini kit (Cat. No. 740609.50) (MACHERY-NAGEL GmbH co.KG, Germany) following the manufacturer‘s instructions. Subsequently, purified DNA was sequenced by Sanger Sequencing (ABIprism 3130) and checked for the specific sequences for each isoform.

### Statistical analyses

Circ-eQTL validation analysis was performed in RStudio 2024.12.1 using R v4.5. To assess the association between genotype and circRNA expression, generalized linear models (GLMs) with a Gamma distribution and identity link function were fitted. The initial model included genotype, sex, group (MS vs HC), and all possible interactions between these factors. As interaction terms did not significantly improve model fit, a simplified model including only the main effects was subsequently evaluated. A Type III analysis of deviance was performed to assess the contribution of each term while adjusting for the others. Where significant genotype effects were detected, post-hoc pairwise comparisons were conducted using estimated marginal means with Tukey’s adjustment for multiple testing. Model comparisons were conducted using likelihood ratio tests and the Akaike Information Criterion (AIC).

In the sQTL validation experiment, for band quantification and circRNA expression analysis, the Kruskal-Wallis test was used to assess differences among genotypes; when significant, Dunn’s post hoc test was applied to determine pairwise group differences.

### Ethics

The Donostia University Hospital Ethics Committee approved the study and all donors provided written informed consent before blood sampling. For the case-control association study, patients provided informed consent at each recruiting center and the institutional ethics committees of these centers approved the study.

## RESULTS

### Circ-eQTL

The circ-eQTL analysis identified 42,077 significant circ-eQTLs (FDR < 0.05 and MAF > 0.1; Supplementary Table 1). Following the intersection with GWAS-reported loci and sQTLs from the GTEx project, two circ-eQTLs were prioritized for validation. The first candidate was rs7214410– hsa_circ_0106983 (FDR = 6.56 × 10^−6^). Additionally, we selected rs11079784, another significant circ-eQTL (FDR = 1.81 × 10^−2^), which not only appeared in our analysis but also represents the lead SNP within the chromosome 17 locus previously associated with multiple sclerosis (MS). The second candidate selected for validation based on our prioritization strategy was rs6498184– hsa_circ_0002161(FDR = 3.39 × 10^−2^)(Table 2).

**Table 2.**
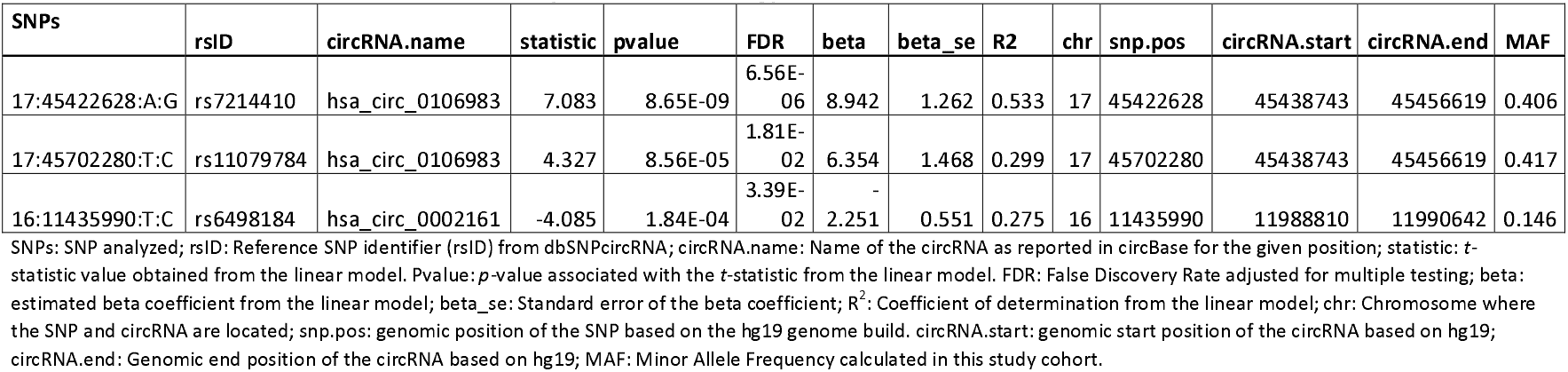
circ-eQTL selected for validation from the prioritization strategy.

**Table 3.**
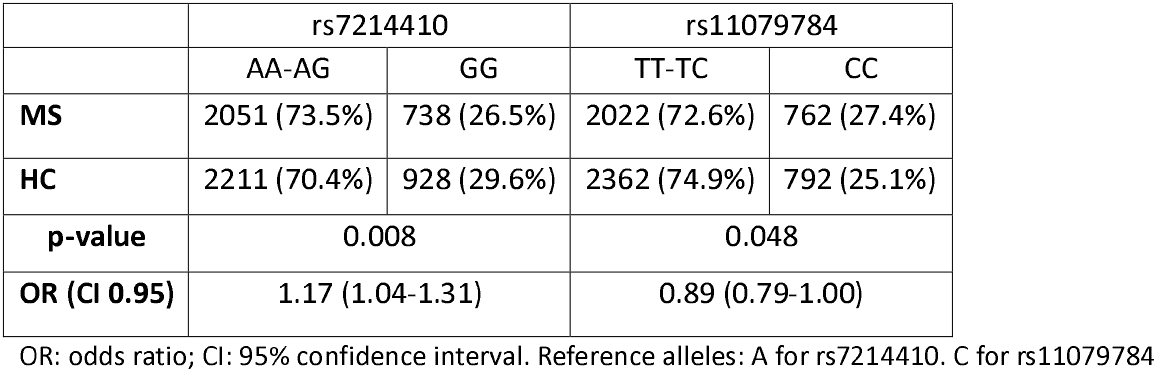
Genotype distributions obtained in the case-control association study are shown according to the dominant model, which showed the best results in the association study.

The validation study of our candidates applying the GLM analysis for rs7214410 revealed significant effects of genotype (p < 0.0001), sex (p = 0.004), and group (p = 0.026) on hsa_circ_0106983 expression levels. Post-hoc pairwise comparisons indicated that expression levels differed significantly among the three genotype groups. Specifically, individuals heterozygous or homozygous for the alternative allele (AG and GG) exhibited lower hsa_circ_0106983 expression compared to those homozygous for the reference allele (AA) (Fig. 2A; Supplementary Table 2). In contrast, the analysis for rs11079784 identified significant effects of genotype (p < 0.0001) and sex (p < 0.001), while no significant effect was observed for the group (p = 0.307). Post-hoc comparisons confirmed significant differences in hsa_circ_0106983 expression across the three genotype groups (Fig. 2B; Supplementary Table 2). Finally, the validation analysis for rs6498184– hsa_circ_0002161 did not reveal any effect of genotype, sex, or group on hsa_circ_0002161 expression (Supplementary Fig. 1; Supplementary Table 2).

**Figure 2.**
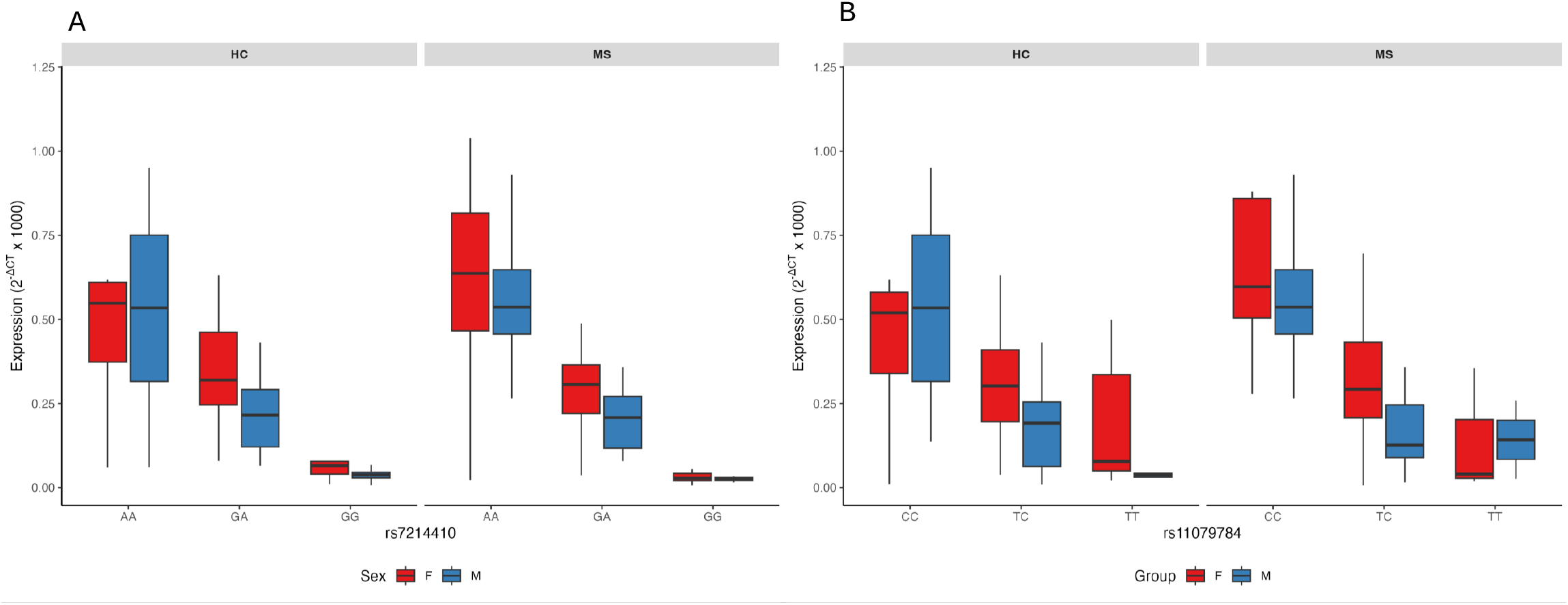
hsa_circ_0106983 circ-eQTL validation. Hsa_circ_0106983 expression is represented according to rs7214410 (A) and rs11079784 (B) genotypes, segregated by sex and group. HC: healthy controls group. MS: multiple sclerosis group. F: female. M: male.

### Case-control association study

The rs7214410 SNP is located in chromosome 17 and colocalizes with an MS-associated region, where the other second circ-eQTL candidate rs11079784 was previously reported as the lead SNP. We wanted to test whether the association of this region with MS could be due to rs7214410 SNP. First, we checked the linkage disequilibrium (LD) between these two variants in the European population, which was of r =0.792. Then, we performed a case-control association study with 2831 PwMS and 3191 HC to see if rs7214410 was associated with the disease and to compare it with the best-associated variant from GWAS (rs11079784). Results shown in Table 2 indicate that rs7214410 shows a stronger association with MS (p=7.9 × 10^-3^; OR= 1.17 (1.04-1.31) than the one that was described before as the lead SNP of the region (p=4.8 × 10^-2^; OR=0.89 (0.79-1.00)(Table 2)).

### Validation of the sQTL

In this study, we identified rs7214410 as a SNP that modulates the expression of hsa_circ_0106983. This variant has been previously reported as a sQTL for the *EFCAB13* gene (187), which, according to the circBase database, is the host gene of hsa_circ_0106983. Thus, we aimed to validate the splicing event in *EFCAB13* by assessing whether exon loss occurred between exons 8 and 11 in relation to the rs7214410 genotype.

The experiment conducted in LCL samples confirmed a genotype-dependent loss of exons. Specifically, individuals homozygous for the G allele lacked the upper band at 397 bp, and the middle band at 253 bp was barely detectable. In contrast, the lower band at 109 bp appeared more intense in these individuals. These findings confirm that exons 9 and 10 are skipped in individuals with the “GG” genotype for rs7214410 (Fig. 3A).

**Figure 3.**
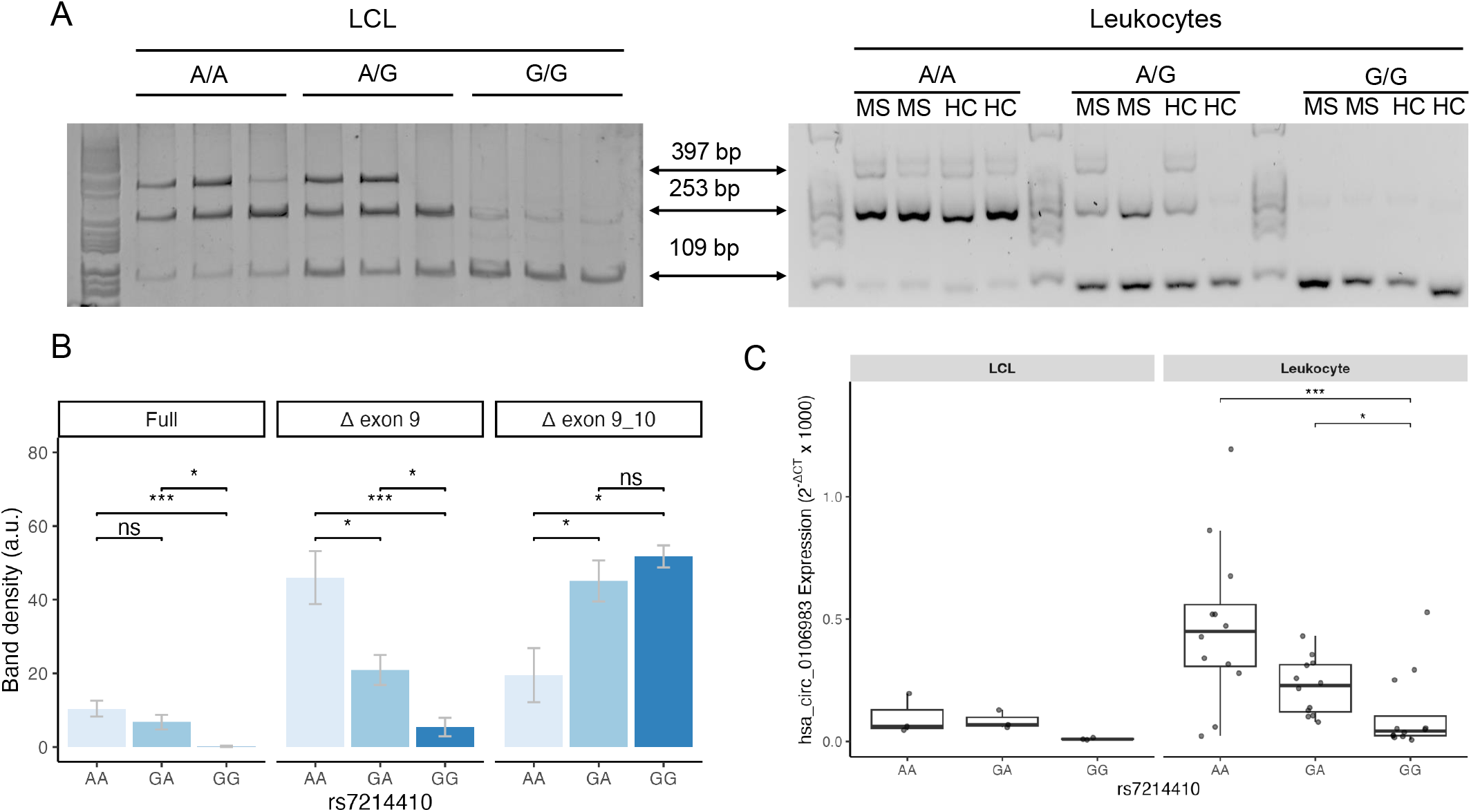
sQTL validation results. A) Gel electrophoresis analysis of the three alternative splicing isoforms of the *EFCAB13* gene. The gel displays bands corresponding to isoforms lacking exon 9 (253 bp band) and both exons 9 and 10 (109 bp band). On the left, LCL samples are shown (n=3 per genotype) and on the right a representative gel from peripheral leukocyte samples (n=2 per genotype) from pwMS and HCs is presented. B) Quantification results of band intensities from leukocyte gel electrophoresis are shown (n=12 per genotype). Blue bars represent the average band density for each genotype, grouped by transcript isoform. C) Expression levels of hsa_circ_010698 are represented grouped by rs7214410 genotype both in LCL and leukocyte samples.

We replicated the experiment using peripheral leukocytes from a subset of pwMS and HC participants of our validation cohort. Consistent with the LCL results, exon loss was observed in heterozygous and “GG” individuals (Fig. 3A). Moreover, band density quantification from electrophoresis gels from leukocyte samples demonstrated that both heterozygous and “GG” homozygous individuals exhibited a significant reduction in the full-length transcript (p=8.3×10^−4^) and in the transcript lacking exon 9 (p=1.7×10^−4^). Conversely, the transcript lacking both exons 9 and 10 showed a significant increase (p=0.03). According to pairwise comparison, the transcript lacking exon 9 has different densities among the three genotypes, while the isoform lacking exons 9 and 10 differs between “AA” homozygous individuals and the other two groups (Fig. 3B).

Additionally, the expression of hsa_circ_0106983 was reduced in “GG” individuals compared to both “AA” (p=0.004) and “GA” genotypes (p=0.03) in peripheral leukocytes, although in LCL samples, these differences did not reach statistical significance (p=0.06) (Fig. 3C). Subsequently, using a linear model, we assessed *EFCAB13* expression levels in the discovery cohort across the three genotypes and observed no significant differences (p=0.2) (Supplementary Fig. 2).

Finally, as a confirmatory experiment, we sequenced the different splicing isoforms. In the isoform retaining all exons, exon 8 was followed by exon 9. In the isoform lacking exon 9, exon 8 was directly followed by exon 10. Lastly, in the isoform in which both exons 9 and 10 were skipped, exon 8 was directly joined to exon 11 (Supplementary Fig. 3).

## DISCUSSION

In recent years, circRNAs emerged as an important player in transcriptome regulation and they have been related to many human diseases, including multiple sclerosis[7],[21]–[23]. Several large-scale studies in the last decade have revealed the regulatory nature of genomic variants increasing our knowledge about gene expression regulation[24],[25]. In addition, some authors have focused on studying eQTLs that affect the expression of miRNAs (miR-eQTLs) or long non-coding RNAs (lncRNAs) [14],[26],[27]. However, to the best of our knowledge, there is only a work in which a comprehensive analysis of circ-eQTL has been done, reporting that circ-eQTLs exist independently of eQTLs[28].

In this work, we have performed a comprehensive circ-eQTL analysis in the context of multiple sclerosis and we have identified 42.077 *cis-*circ-eQTLs in peripheral blood leukocytes isolated from people with MS and healthy individuals. To select the most relevant candidates, we have intersected the list of circ-eQTL with variants associated with MS risk, as well as with previously described sQTL. Our approach gave us two candidates, from which the rs7214410-hsa_circ_0106983 association was confirmed. In addition, this SNP has been also identified as an sQTL that affects the splicing of *EFCAB13*.

To date, there is only another work studying circ-eQTL in multiple sclerosis[16]. Cardamone and colleagues, performed an eQTL analysis to search for association between circRNA expression and genotype and found 89 circ-eQTL. The number of circ-eQTL in our study is much higher (42.077) but this could be due to several methodological differences between the two studies, such as sample size or the approach used for circ-eQTL identification. Cardamone et al., selected the variants to include in circ-eQTL analysis based on MS GWAS that had previously identified SNPs associated with the disease. Nonetheless, our approach explored all the possible circ-eQTLs and then focused on those that lay in GWAS-associated regions to select the most relevant associations. Interestingly, Cardamone et al. identified rs11079784 SNP as a circ-eQTL, but the circRNA associated was hsa_circ_0000777, which is in the same region as hsa_circ_0106983[16]. Both studies support the idea that in this chromosome 17 region, which has been previously associated with the MS risk, there are SNPs altering circRNA expression. However, probably due to differences in selecting putative circ-eQTL candidates, the SNP evidenced as eQTL for circRNA is different in both studies.

The host gene of these circRNAs is *EFCAB13* (EF-hand calcium-binding domain-containing protein 13), a protein that contains a calcium-binding domain that is shared by a variety of calcium sensor proteins. It exhibits low expression across many tissues, with the highest expression levels observed in testis or thyroid gland[25]. Its function remains unknown, although it has been suggested to play a role in neuronal function and plasticity[29],[30]. Interestingly, our data reveal that the SNP causing the circ-eQTL (rs7214410) is also an sQTL for *EFCAB13* gene. Of note, other authors have identified a different SNP in the same region (rs3851808) as an sQTL affecting the alternative splicing in *EFCAB13*[31]. Both rs3851808 and rs7214410 show LD (r^2^= 0.706 in the European population CEU); however, according to our results, rs3851808 is not a circ-eQTL and therefore, it was not considered as a candidate for colocalization with sQTL data. In addition, candidate SNP prioritization used by Putscher et al. was knowledge-based, starting with MS-risk SNPs. In contrast, we carried out the analysis prior to focusing on MS-associated regions. Furthermore, Putscher et al. used RNA isolated from B cells whereas in our study we analyzed RNA from the overall leukocyte population, which might affect the sQTL effect[25]. Moreover, low-frequency variants in *EFCAB13* have also been associated with MS susceptibility in an Italian population[32], highlighting this region as a candidate for further investigation.

Taking into account that circRNA biogenesis can be considered a form of alternative splicing, it is very interesting to observe that the same variant can affect both alternative splicing of *EFCAB13* and the expression of one of the circRNAs encoded in the same region. Of note, we found that the presence of the alternative allele (G) of rs7214410 decreases the expression of hsa_circ_0106983 and increases the isoform lacking exons 9 and 10. Yet, this variant does not affect the expression of the linear transcript, and previous studies have also reported no significant differences in expression levels of *EFCAB13* between MS and HC[32]. Whether these events are related to each other or there are independent events has still to be determined. One possible hypothesis is that in the presence of G allele, the circRNA biogenesis machinery favors the synthesis of other circRNA that include exon 9 and 10. Overall, our data suggest that variants in this region may exert functional effects primarily on non-coding RNAs, rather than on coding transcripts.

The case-control association study carried out in this work has revealed that rs7214410 is strongly associated with MS, suggesting that this variant could be the causal SNP in this region, instead of the GWAS-reported SNP (rs11079784). Our data point to the importance of focusing also on the non-coding part of the genome as causal of disease association. The function of this circRNA and how this variation in its expression could affect leukocyte function deserves further investigation. All data suggest that we should keep investigating in this direction to uncover the relation of this region with MS susceptibility. Evidence gathered by several groups point at this region around *EFCAB13* to have a role in MS-susceptibility, having a functional effect on splicing and/or expression of nearby genes or circRNAs.

## ACKNOWLEDGEMENTS

The authors thank all the participants in the study.

## AUTHOR CONTRIBUTIONS

**Saioa Gs Iñiguez**: investigation, formal analysis, validation, visualization, writing – original draft. **Leire Iparraguirre**: investigation, formal analysis. **Eduardo Andrés-León**: formal analysis. **Hirune Crespillo**: investigation, resources. **Leire Romarate**: resources. **Tamara Castillo Triviño:** resources. **Elena Urcelay, Manuel Comabella, Sunny Malhotra, Xavier Montalban, Lluís Ramió-Torrentà, Anna Quiroga-Varela, Koen Vandenbroeck, Ane Aldekoa, Antonio Alcina:** resources, validation. **David Otaegui:** conceptualization, formal analysis, funding acquisition, project administration, supervision, resources. **Fuencisla Matesanz:** formal analysis, investigation, resources, validation. **Maider Muñoz-Culla:** conceptualization, formal analysis, funding acquisition, project administration, supervision, visualization, Writing – original draft. **All the authors:** Writing – review & editing.

## Declaration of generative AI and AI-assisted technologies in the writing process

During the preparation of this work the authors used ChatGPT Open AI to improve English and readability of the manuscript. After that, the authors reviewed and edited the content as needed and take full responsibility for the content of the published article.

---

**Supplementary figure 1.**
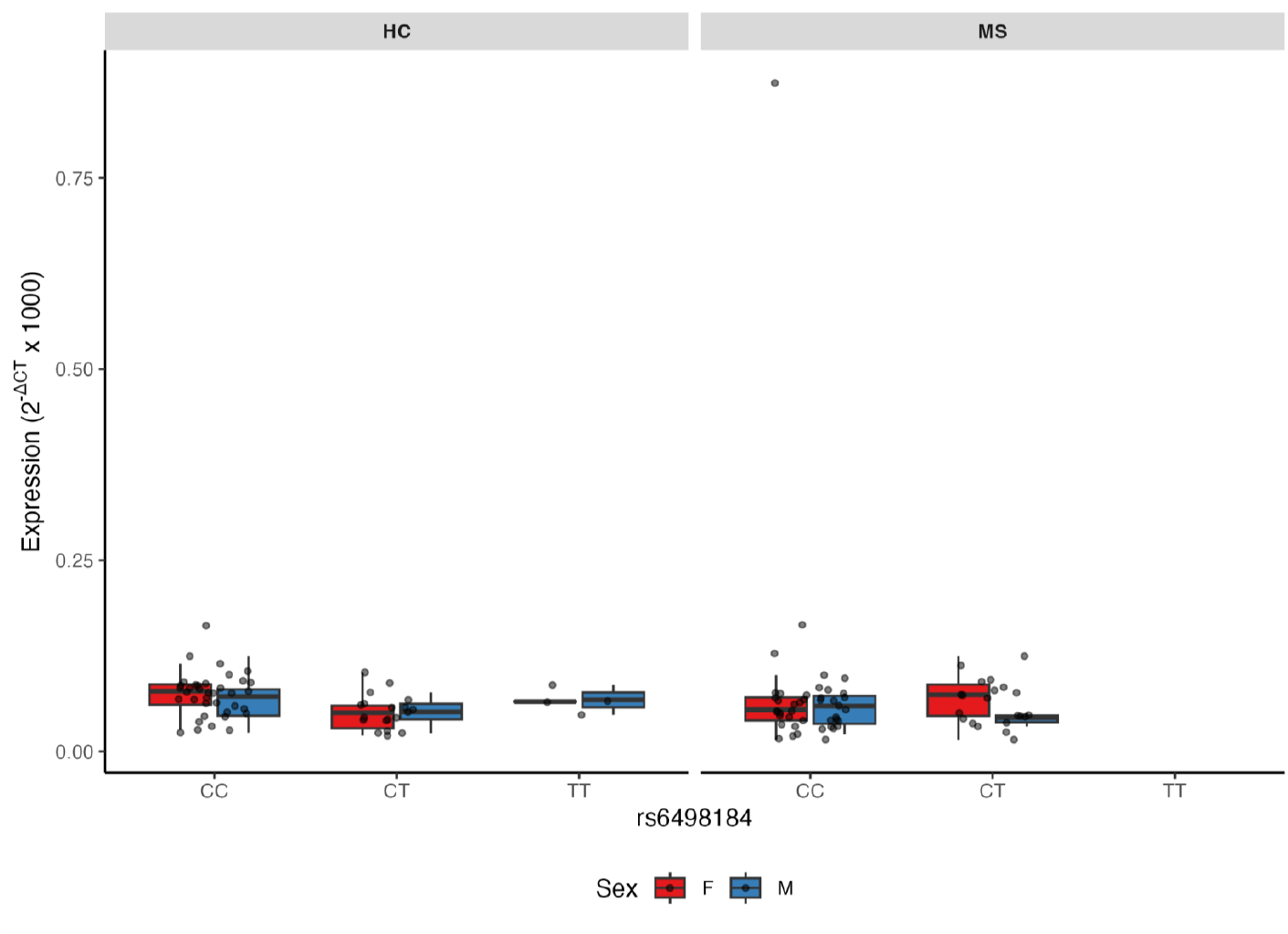
hsa_circ_0002161 circ-eQTL validation. Hsa_circ_0002161 expression according to rs6498184 genotype, segregated by sex and group. HC: healthy controls group. MS: multiple sclerosis group. F: female. M: male.

**Supplementary figure 2.**
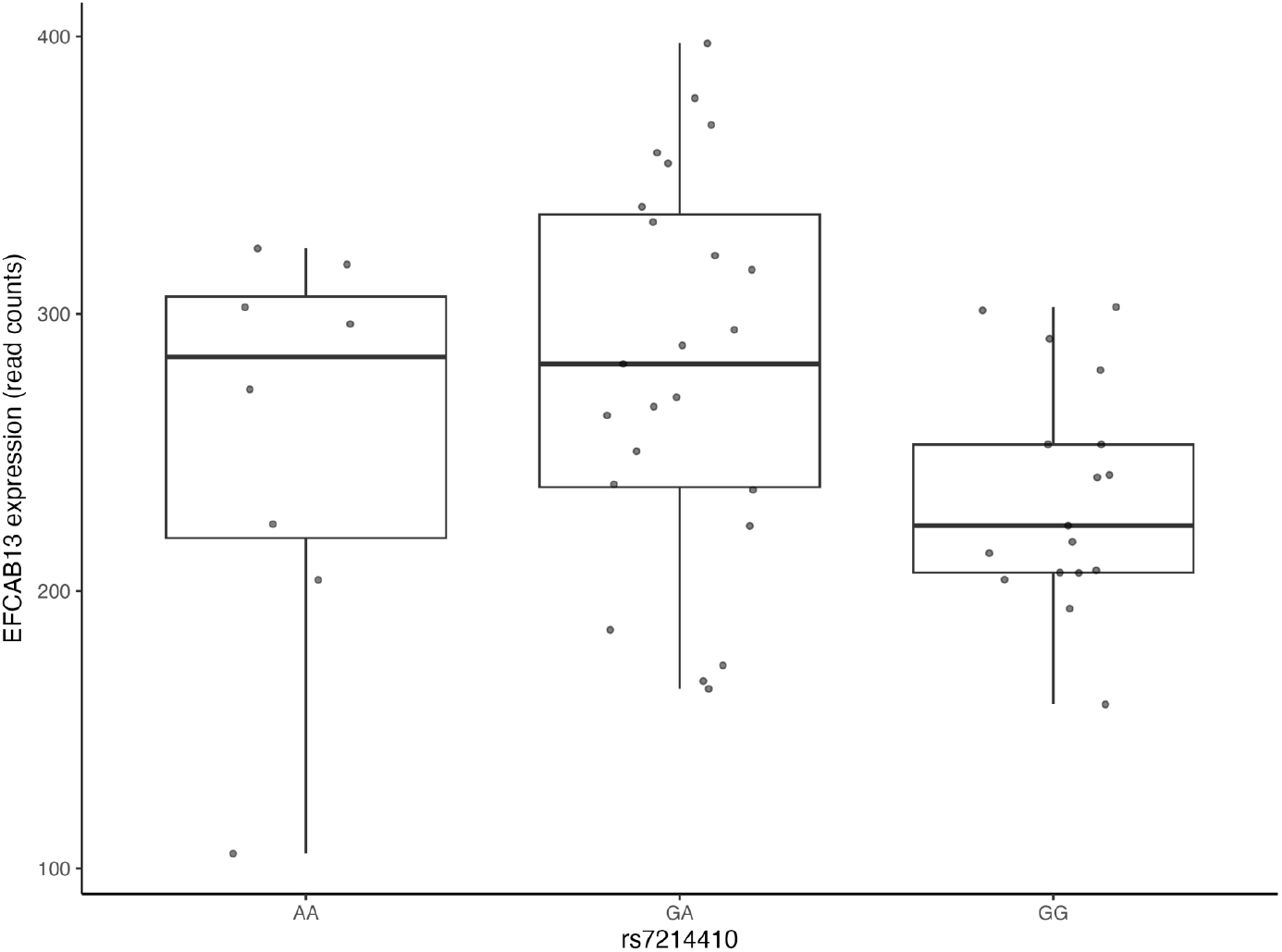
EFCAB13 expression (normalized read counts) in the dicovery cohort samples according to rs7214410

**Suplementary figure 3.**
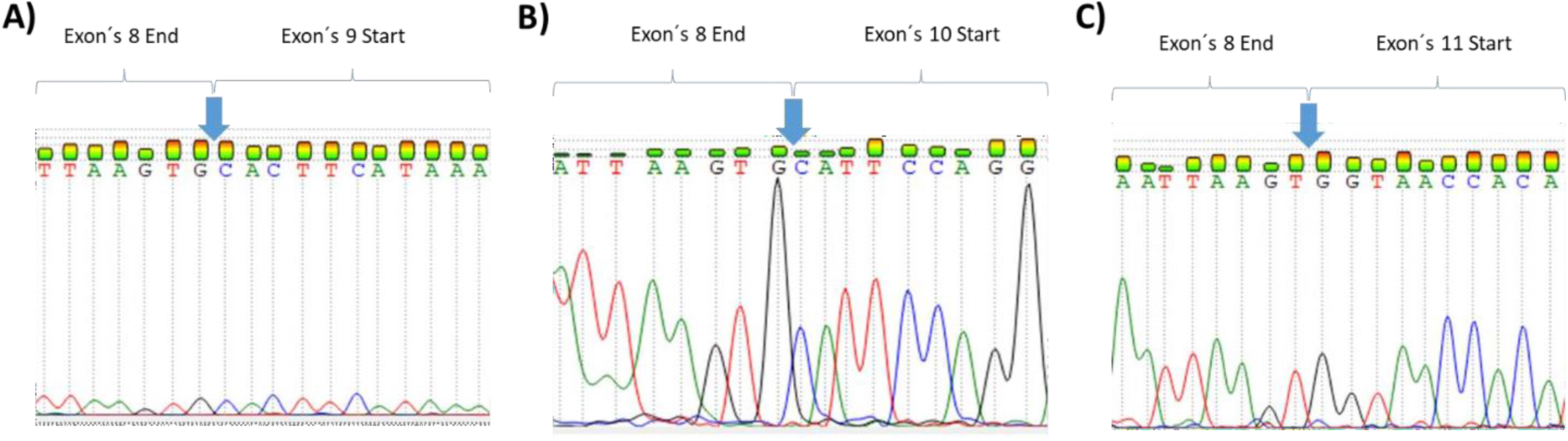
Representation of the different splicing isoforms sequences. A) Isoform retaining all exons. B) Isoform skipping exon 9. C) Isoform skipping both exon 9 and 10.

