## Supplementary table 2 for "A dual-function variant on chromosome 17 regulates circRNA expression and splicing in multiple sclerosis"

Supplementary table 2. circ-eQTL validation analysis results.

| **rs7214410** | | | | |
| --- | --- | --- | --- | --- |
| ***Generalized linear model coefficients*** | | | | |
| **Predictor** | **Estimate** | **Std. Error** | **t-value** | **p-value** |
| (Intercept) | 0.615 | 0.0940 | 6.540 | 1.39e-09 *** |
| Genotype GA vs AA | -0.272 | 0.0989 | -2.753 | 0.00679 ** |
| Genotype GG vs AA | -0.495 | 0.0941 | -5.265 | 5.86e-07 *** |
| Sex male (vs female) | -0.0566 | 0.0180 | -3.135 | 0.00214 ** |
| Group MS (vs HC) | -0.0410 | 0.0175 | -2.340 | 0.02088 * |
| ***Type III Analysis of Deviance (Likelihood Ratio Tests)*** | | | | |
| **Factor** | **LR Chisq** | **Df** | **p-value** |  |
| Genotype rs7214410 | 82.022 | 2 | < 2.2e-16 *** | |
| Sex | 8.119 | 1 | 0.00438 ** |  |
| Group (MS vs HC) | 4.969 | 1 | 0.02580 * |  |
| ***Post-hoc pairwise comparisons (Tukey adjustment)*** | | | | |
| **Contrast** | **Estimate** | **Std. Error** | **t-value** | **p-value** |
| AA vs GA | 0.272 | 0.0989 | 2.753 | 0.0185 |
| AA vs GG | 0.495 | 0.0941 | 5.265 | < 0.0001 |
| GA vs GG | 0.223 | 0.0342 | 6.530 | < 0.0001 |
| **rs11079784** | | | | |
| ***Generalized linear model coefficients*** | | | | |
| **Predictor** | **Estimate** | **Std. Error** | **t-value** | **p-value** |
| (Intercept) | 0.592 | 0.0941 | 6.288 | 4.91e-09 *** |
| Genotype TC vs CC | -0.285 | 0.0954 | -2.984 | 0.00343 ** |
| Genotype TT vs CC | -0.404 | 0.0937 | -4.311 | 3.27e-05 *** |
| Sex male (vs female) | -0.140 | 0.0323 | -4.333 | 2.99e-05 *** |
| Group MS (vs HC) | 0.0291 | 0.0322 | 0.905 | 0.3673 |
| ***Type III Analysis of Deviance (Likelihood Ratio Tests)*** | | | | |
| **Factor** | **LR Chisq** | **Df** | **p-value** |  |
| Genotype rs11079784 | 37.805 | 2 | 6.18e-09 *** |  |
| Sex | 14.368 | 1 | 0.0001503 *** | |
| Group (MS vs HC) | 1.042 | 1 | 0.3074 |  |
| ***Post-hoc pairwise comparisons (Tukey adjustment)*** | | | | |
| **Contrast** | **Estimate** | **Std. Error** | **t-value** | **p-value** |
| CC vs TC | 0.285 | 0.0954 | 2.984 | 0.0095 |
| CC vs TT | 0.404 | 0.0937 | 4.311 | 0.0001 |
| TC vs TT | 0.119 | 0.0320 | 3.729 | 0.0008 |
| **rs6498184** | | | | |
| ***Generalized linear model coefficients*** | | | | |
| **Predictor** | **Estimate** | **Std. Error** | **t-value** | **p-value** |
| (Intercept) | 0.0771 | 0.0115 | 6.684 | 7.93e-10 *** |
| Genotype CT vs CC | -0.0172 | 0.0124 | -1.385 | 0.169 |
| Genotype TT vs CC | -0.0036 | 0.0352 | -0.103 | 0.918 |
| Sex male (vs female) | -0.0120 | 0.0126 | -0.953 | 0.343 |
| Group MS (vs HC) | 0.00314 | 0.0123 | 0.254 | 0.800 |
| ***Type III Analysis of Deviance (Likelihood Ratio Tests)*** | | | | |
| **Factor** | **LR Chisq** | **Df** | **p-value** |  |
| Genotype rs6498184 | 1.786 | 2 | 0.4094 |  |
| Sex | 0.783 | 1 | 0.3762 |  |
| Group (MS vs HC) | 0.0595 | 1 | 0.8072 |  |
| ***Post-hoc pairwise comparisons (Tukey adjustment)*** | | | | |
| **Contrast** | **Estimate** | **Std. Error** | **t-value** | **p-value** |
| CC vs CT | 0.0172 | 0.0124 | 1.385 | 0.3522 |
| CC vs TT | 0.00362 | 0.0352 | 0.103 | 0.9942 |
| CT vs TT | -0.0136 | 0.0351 | -0.387 | 0.9209 |
